# Optimization of a maize rapid cycle breeding scheme using the Modular Breeding Program Simulator (MoBPS)

**DOI:** 10.1101/2025.01.10.632416

**Authors:** Torsten Pook, Mila Leonie Tost, Henner Simianer

**Affiliations:** Department of Animal Sciences, Animal Breeding and Genetics Group, University of Goettingen, 37075 Goettingen, Germany; Center for Integrated Breeding Research, University of Goettingen, 37075 Goettingen, Germany; Animal Breeding and Genomics, Wageningen University & Research, 6700 AH Wageningen, The Netherlands; Department of Crop Sciences, Plant Breeding and Methodology Group, University of Goettingen, 37075 Goettingen, Germany

**Keywords:** rapid cycle, breeding, simulation, maize, speed breeding, rapid generation advance, plantae

## Abstract

In recent years, the turnover of plant breeding has substantially increased as the use of genomic information allows for earlier selection and the integration of controlled growing environments reduces time to reach a particular growing stage. However, high generation turnover and intensive selection of lines before own yield trials are performed come at the risk of a drastic reduction of genetic diversity paired with lower prediction accuracies. To this end, we investigate strategies to cope with these challenges in a maize rapid cycle breeding scheme using stochastic simulations using the software MoBPS. We find that genetic gains soon reach a plateau when only the original breeding material is phenotyped. Updating the training data set via additional phenotyping of crosses or doubled haploid lines ensures long-term progress with a gain of 6.80 / 6.95 genetic standard deviations for the performance as a cross / *per se* after 30 cycles of breeding compared to 3.40 / 4.28 without additional phenotyping. Adding genetic material with comparable genetic level and novel diversity from outside the breeding material led to a further increase to 9.34 / 7.89 genetic standard deviations. In particular, for the management of genetic diversity, further additions to the breeding scheme are analyzed to optimize the number of selected lines per cycle and to account for the relatedness of F2 plants in the selection using the software AlphaMate. Finding a balance between genetic gains and diversity is important for a given time frame. MoBPS provides a tool for the quantification of these effects and provides solutions specific to the respective breeding program.

## Introduction

Plant breeding is a complex task with the ultimate goal of producing crop varieties with unique and superior traits, used in human and animal nutrition or as an energy source for green energy. Breeders are trying to improve the characteristics of varieties continuously and thereby achieve sustainable and long-term genetic gain. With rising demands for higher production levels and more complex breeding goals, plant breeders are permanently challenged to enhance breeding strategies and efforts (Foley *et al*. 2011).

Genetic gain can be increased by controlled mating, which leads to an increase in heritability, higher selection intensity, or reduced time between generations (Falconer and Mackay 1996). The options for further increasing the selection intensity are limited, as this might reduce genetic diversity (Falconer and Mackay 1996). The generation interval, and thus the capability to perform several breeding cycles within a certain time frame, is an important factor impacting genetic gain (Falconer and Mackay 1996). Consequently, large-scale breeding companies often ship breeding material between the northern and southern hemispheres to achieve two generation cycles per year. By the use of the controlled environment of a greenhouse with supplemental lighting, heating, and advanced modern crop breeding technologies the number of cycles per year can be increased even further, with speed-breeding approaches enabling up to six cycles for wheat, barley, and chickpea, or four cycles per year in canola (Watson *et al*. 2018; Ghosh *et al*. 2018). With the introduction of genomic selection (Meuwissen *et al*. 2001; Schaeffer 2006; Jannink 2010; Heslot *et al*. 2015) this process can be made even more efficient as reliable breeding value are available earlier and lines can be selected before own phenotypes are available, with approaches such as rapid cycle breeding (Gerald *et al*. 2013; Crossa *et al*. 2017; Gorjanc and Hickey 2018). Thus, rapid cycling paired with speed breeding has the potential to be a highly efficient framework for various applications in plant breeding like variety development, pre-breeding, and bridging efforts to introduce new genetic material into an existing breeding program (Gorjanc *et al*. 2016; Allier *et al*. 2020). In recent years, pre-breeding has become increasingly relevant for the utilization of genetic resources and for broadening the genetic basis of elite germplasm (Smith *et al*. 2015; Gaikpa *et al*. 2021). In this sense, pre-breeding can be a tool to make material of high genetic diversity useable for breeding (Mayer *et al*. 2018, 2020). However, as these concepts have only recently been developed, practical yield trials and studies have only considered the use of these breeding scheme for a couple of breeding cycles (Dreisigacker *et al*. 2023; Polzer 2023). However, as modern varieties are much superior for performance traits like yield compared to more diverse landrace material or old breeding populations (Voss-Fels *et al*. 2019), it will be essential to pursue a rapid cycling breeding scheme for many generations to bridge the gap between exotic and elite breeding material.

This in turn comes with various challenges (Wanga *et al*. 2021) - particularly in the context of phenotyping. Phenotyping of some traits is only possible in adult plants and/or requires seed propagation steps (Watson *et al*. 2018). Thus, phenotypic data is less reliable and/or only available for plants that are already several cycles behind the current breeding material. This raises new questions such as whether and when genomic prediction models will break down as they are only supplied with phenotypic data from past breeding cycles, hence leading to lower prediction accuracies (Lorenz 2013; Neyhart *et al*. 2017). Additionally, genetic diversity might decrease with a high number of breeding cycles and further reduced prediction accuracy (Sallam *et al*. 2015; Schopp *et al*. 2017). Therefore, a detailed consideration concerning the management of genetic diversity and inbreeding is needed (Smith *et al*. 2015; Louwaars 2018).

Although quantitative genetic theory provides indications of the expected effect of breeding actions, changing one parameter in a breeding program inevitably affect other parameters. Hence, modern breeding programs are usually too complex to reliably assess the effect of a single isolated breeding decision (Pook *et al*. 2021), in particular if the relative effects are small compared to the underlying stochasticity in a breeding scheme. Therefore, breeding decisions often rely on the breeder’s experience (Duvick 2002) that is often based on previous years and a limited number of real-world breeding experiments. As financial resources are limited, computer simulations (Sargolzaei and Schenkel 2009; Pook *et al*. 2020b; Gaynor *et al*. 2021; Villiers *et al*. 2022) can be a useful tool to aid the decision-making process in a breeding program and complement real-world breeding experiments.

Due to the increase in computational power in recent years, the use of stochastic simulations for quantitatively evaluating breeding programs using tools such as MoBPS (Pook *et al*. 2020b) has been suggested with various examples from practical breeding programs in animal breeding (Büttgen *et al*. 2020; Pook *et al*. 2021; Martin *et al*. 2023; Hassanpour *et al*. 2023). Although the MoBPS framework has all the required features for the realistic modelling of plant breeding programs, there are, to date, no published exemplary applications of the framework for complex plant breeding programs.

An important component of long-term breeding is the management of genetic diversity with approaches like optimum genetic contribution selection (Meuwissen 1997), which is being widely used, particularly in animal breeding. However, direct application in plant breeding is difficult, as it is for example common practice to work with a limited number of selected plants with equal contributions. To handle these kinds of restrictions, approaches such as AlphaMate (Gorjanc *et al*. 2018) and MateSel (Kinghorn 2011) have been proposed which suggest mating plans under a set of predefined conditions and optimize these via evolutionary algorithms, instead of plainly setting contribution levels of each potential parent under the constrain of a set inbreeding rate (Meuwissen 1997). Beyond this, more attention has been given to not only considering the expected performance of an offspring but also its variance, with approaches that consider Mendelian sampling variance (Bonk *et al*. 2016; Allier *et al*. 2019; Bijma *et al*. 2020; am Niehoff *et al*. 2024)

In this work, we evaluate the potential of speed breeding exemplified by a rapid cycle breeding scheme in maize (*Zea mays*) by employing stochastic simulations in MoBPS (Pook *et al*. 2020b). A particular focus of the analysis is on the management of genetic diversity through selection and introduction of new material with comparable genetic level and novel diversity from outside of the breeding pool over a longer time horizon. Furthermore, differences between direct selection on traits phenotyped in crosses compared to indirect selection by only phenotyping DH lines *per se* are analyzed.

## Materials and methods

As a baseline scenario, we considered a breeding scheme inspired by the rapid cycle breeding scheme from the MAZE project (http://www.europeanmaize.net/) as first suggested by Polzer (2023), aiming to use pre-breeding to enhance performance from material of a European maize landrace. In the project, which is not directly transferable to a commercial breeding program, only three plant height (PH) traits at different growth stages (V4, V6, final) were considered for selection; for details on the different growing stages see Hölker *et al*. (2019). The heritabilities and genetic correlations between the traits were taken from Hölker *et al*. (2019). In MoBPS, the three traits were simulated in two variants, one representing the performance as an inbred line *per se* and the other performance as a test cross, hereafter referred to as *per se* and cross performance. Correlations between the performance of the same trait when phenotyped *per se* and as a cross were taken from Hölker *et al*. (2019) while remaining genetic and residual correlations were meaningfully completed as shown in Table 1. Whenever phenotyping was performed, the corresponding version (*per se* / cross) of the trait was phenotyped. Each trait was simulated to be quantitative with 500 purely additive QTLs, 500 purely dominant QTLs, 100 qualitative epistatic QTLs, and 100 quantitative epistatic QTLs each (n.additive, n.dominant, n.qualitative, n.quantitative in MoBPS (Pook *et al*. 2020b)). Due to the genetic correlation between traits, QTLs of each trait also impact the other traits. The interested reader is referred to MoBPS User Guidelines at https://github.com/tpook92/MoBPS and for details on the modelling of trait correlations in MoBPS.

**Table 1.** Heritabilites (main diagonal), Genetic correlations (above diagonal), and residual correlation (below diagonal) of the simulated traits.

| Trait | DH |  |  | cross |  |  |
| --- | --- | --- | --- | --- | --- | --- |
|  | PH_v4 | PH_v6 | PH_final | PH_v4 | PH_v6 | PH_final |
| PH_v4 (DH) | 0.95 | 0.94 | 0.50 | 0.62 | 0.56* | 0.20* |
| PH_v6 (DH) | 0.30 | 0.95 | 0.58 | 0.61* | 0.68 | 0.30* |
| PH_final (DH) | 0.10 | 0.15 | 0.96 | 0.25* | 0.34* | 0.78 |
| PH_v4 (cross) | 0.20 | 0.06 | 0.02 | 0.76 | 0.86 | 0.21 |
| PH_v6 (cross) | 0.06 | 0.20 | 0.03 | 0.30 | 0.77 | 0.33 |
| PH_final (cross) | 0.02 | 0.03 | 0.20 | 0.10 | 0.15 | 0.87 |

Genetic correlations that were meaningfully completed are indicated with \*.

A schematic overview of the breeding program is given in Figure 1. The breeding program as implemented in the MoBPS interactive environment from Pook *et al*. (2020a) is given in Supplementary Figure S1. As a starting point of the breeding program, an initial pool of 409 founder DH lines of a European maize landrace was used (Mayer *et al*. 2020). Initially, 10 DHs were selected based on performance in the selection index with breeding values estimated via genomic best linear unbiased prediction (gBLUP) (Meuwissen *et al*. 2001) with phenotypes being simulated for the initial DHs. Subsequently, selected DHs were combined in a half-diallelic crossing scheme to generate offspring from all 45 possible genotype combinations. Crosses were subsequently selfed to generate 23 offspring from each combination, resulting in a pool of 1035 F2 lines in cycle 1 (C1F2). From these, the best performing 30 C1F2 were selected based on a breeding value estimation based on the phenotypes from original DHs lines. From the top 30 lines, 15 crossing pairs were created at random and 69 offspring C2F2 lines were generated from each mating pair, resulting again in a pool of 1035 lines. To mimic the generation of a final product of the breeding scheme, DH lines were produced from the top three F2 lines. To avoid a severe loss in genetic diversity, at most 10 C1F2 individuals that originated from the same initial DH line were selected in the first cycle. The R-package rrBLUP was used for the estimation of breeding values (Endelman 2011).

**Figure 1.**
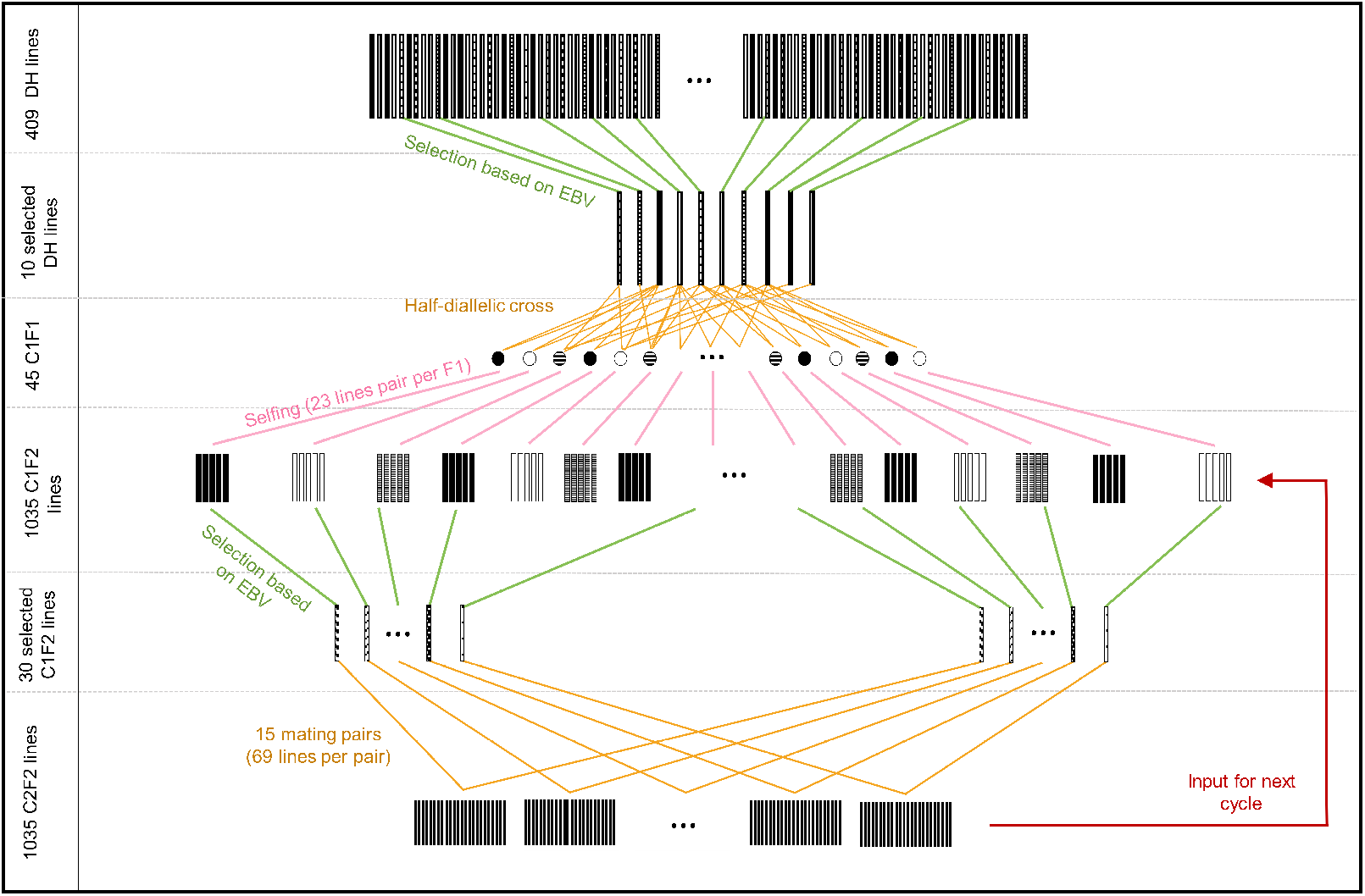
Schematic overview of a rapid cycle breeding scheme.

In contrast to the real-world breeding program in MAZE (Polzer 2023) where only 3 breeding cycles were considered, a considerably longer horizon of 30 breeding cycles was considered in this study. Additionally, the selection of lines was done based on a selection index with equal weight on three traits, aiming to increase the first two traits and to decrease PH_final. In contrast, the real-world breeding scheme used the absolute difference from the current average plant height for PH_final with twice the weight instead of direct selection forqa low final plant height (Polzer 2023). This was done to avoid the effects of a long-term overall change in PH_final affecting the design of the index. In the following, we report only the results concerning on the overall selection index.

In addition to the baseline scenario, various modifications of the breeding system were considered and are described in the following subsections. For all subsequently described scenarios, visualizations based on MoBPSweb (Pook *et al*. 2020a) can be found in the supplementary material (Figure S2 - S4).

### Additional phenotyping

Firstly, additional phenotyping of crosses (Figure S2) was considered. For simplicity, we did not simulate a second gene pool to mimic the mating to a tester and phenotyping as a hybrid. Instead were simulated phenotypes based on the F2 genotypes directly with the appropriate heritability for crosses taken from Hölker *et al*. (2019). Since crosses are phenotyped in a plot and the same tester plant is commonly used for all lines this should not affect the results here. Since this requires an additional seed propagation step in practice, it was assumed that phenotypes for the selected F2 lines are available from two cycles ago. As a second strategy, the generation of DH lines from the F2 lines was considered with a subsequent phenotypic *per se* (Figure S3). Given the cost of DH production and phenotyping, along with the expected technical failure rate (Chaikam *et al*. 2012; Melchinger *et al*. 2016), phenotypes were assumed to only be available for 200 of the 1035 F2 lines. Since DH production in practice requires steps of induction, selfing, multiplication, and phenotyping it was assumed that phenotypes are only available for the lines from four cycles ago. The training population for the breeding value estimation included the last four generations of DHs with available phenotypes, resulting in a training population of 800 lines with *per se* phenotypes compared to the 1035 phenotypes from crosses.

### Genetic diversity

For the management of genetic diversity, three strategies were considered: Firstly, the introduction of new genetic resources from outside the breeding material was considered. For this, new genetic material was simulated in each cycle by using the founder DH lines as a starting point and applying a mild selection intensity in the recurrent selection (200 selected lines from a pool of 409 lines; Figure S4). The recurrent selection was continued until the average genetic value of the population exceeded the average genetic value in the F2 lines, with the index weights being adapted in each cycle to obtain similar genetic progress in each respective trait. Subsequently, material from the previous generation with slightly lower performance was used to replace the selected F2 lines in some of the crosses to generate the F2s of the next cycle. As a baseline, 20% of all matings used external material with various other proportions being considered (10%, 30%, 50%, 70%, 100%). For the breeding value estimation, 200 DH lines from the newly introduced material were generated and phenotyped. To maintain a constant training population size, the last two cycles of DH lines with available phenotypes from within the breeding program and DH lines from the last two newly introduced cohorts were considered, resulting in a training population of 800 lines with *per se* phenotypes. For scenarios where selection is based on phenotypes from crosses, no additional phenotypes from the newly introduced cohorts were generated.

Secondly, the software AlphaMate (Gorjanc *et al*. 2018) was used to identify the 15 required mating pairs. For this, each F2 line was limited to be used at most once (LimitContribtionsMax = 1) and the angle parameter in AlphaMate was used to balance between genetic gain and inbreeding. A low angle in AlphaMate puts a high weight on genetic gain, while a high angle will put more focus on preserving genetic diversity (TargetDegrees = 10, 20, 30, 40, 50, 60, 70).

Lastly, the applied selection pressure was varied. For this, the number of selected lines was varied at two stages in the breeding scheme. Firstly, the number of originally selected DH lines from the initial founder set of 409 lines as the source of initial genetic variation of the population was varied from 5 to 50 (baseline: 10). Secondly, the number of selected F2s per cycle was varied between 10 to 200 with in total 1035 lines to select from (baseline: 30).

### Evaluation metrics

For the evaluation, various metrics were used. The resulting genetic gain for the DH traits represents the genetic gain compared to the initial DH populations. The genetic gain as a cross is derived by comparison of F2 lines to crosses that were generated from the initial unselected DH lines. For the evaluation of the remaining genetic diversity, the variances in the true underlying genomic values within a cohort are calculated. Additionally, the share of heterozygous variants and the share of fixed markers in the F2s were computed. To evaluate the performance of the breeding value estimation, the correlation between the underlying true genomic values and the estimated breeding value, hereafter referred to as the prediction accuracy, is used.

All reported results are averages of 250 simulations per scenario.

## Results

In the baseline rapid cycle scheme, the average genomic value of the F2 lines after 30 cycles of breeding is increased by 3.40 genetic standard deviations (gSD). The average genomic value of the top DH lines is increased by 4.28 gSD compared to the initial DH lines (Figure 2.A).

**Figure 2.**
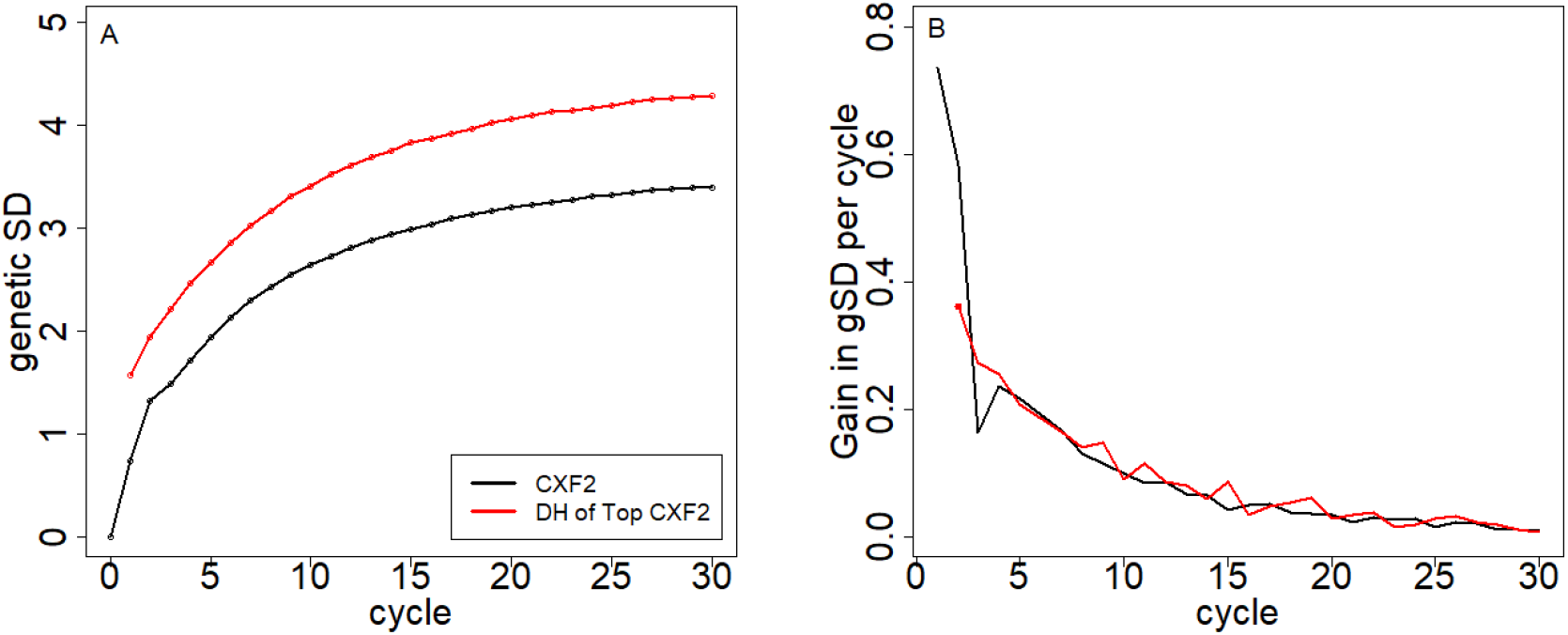
Genetic gain across the breeding cycles (A) and per cycle (B) within the F2 lines and DHs derived from the best F2s. The comparison of performance is against DH lines from the initial founder population and randomly generated crosses from the founders, respectively.

By far the highest gains are obtained in the first two cycles of the breeding scheme. Genetic gains for traits expressed as a cross in cycle 3 are even slightly smaller than in subsequent cycles. After cycle 4 gains are slowly decreasing from around 0.25 gSD per cycle to 0.02 gSD per cycle in cycle 30 (Figure 2.B). Firstly, this is caused by a severe loss in genetic diversity as the genetic standard deviation after 30 cycles is reduced by 74% compared to the initial population (Figure 3.C). Secondly, the genetic distance from the training population with available phenotypes is increasing with a reduced average prediction accuracy of 0.18 in cycle 30 compared to 0.72 in cycle 1 (Figure S5). Especially for the performance in crosses, a plateau is reached after a couple of cycles with basically no further gain with indirect selection on *per se* phenotypes (Figure 2). The share of heterozygous genotypes reduces from 10.9% to 1.6% with 94.3% of all markers being fixed after 30 generations (Table S2 and S3).

**Figure 3.**
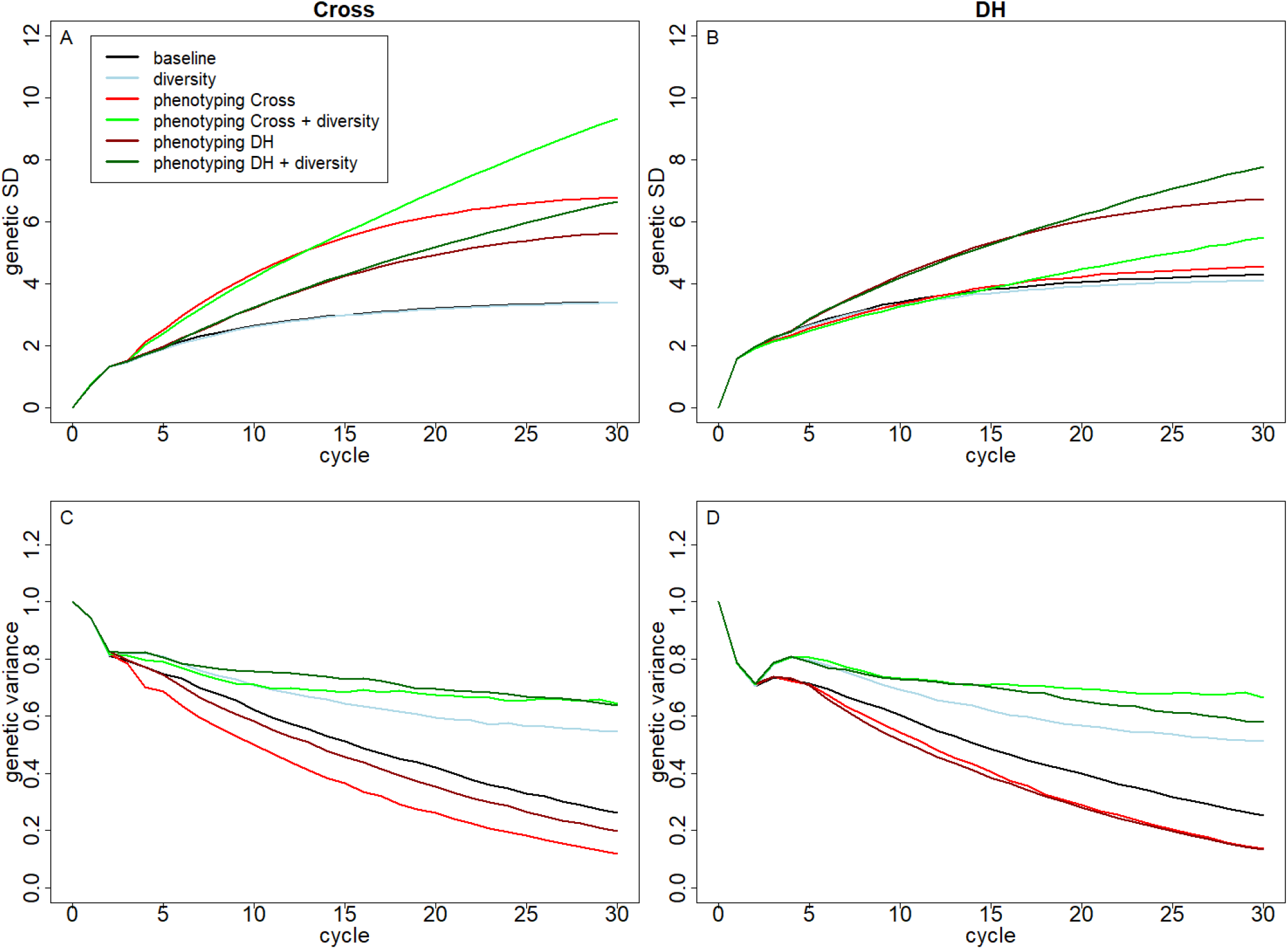
Genetic gain on the selection index for the crosses (A) and DHs (B), as well as the remaining genetic variance of the crosses (C) and DHs (D) in the F2 lines of the respective cycle based on the chosen phenotyping strategy and action against inbreeding.

**Figure 4.**
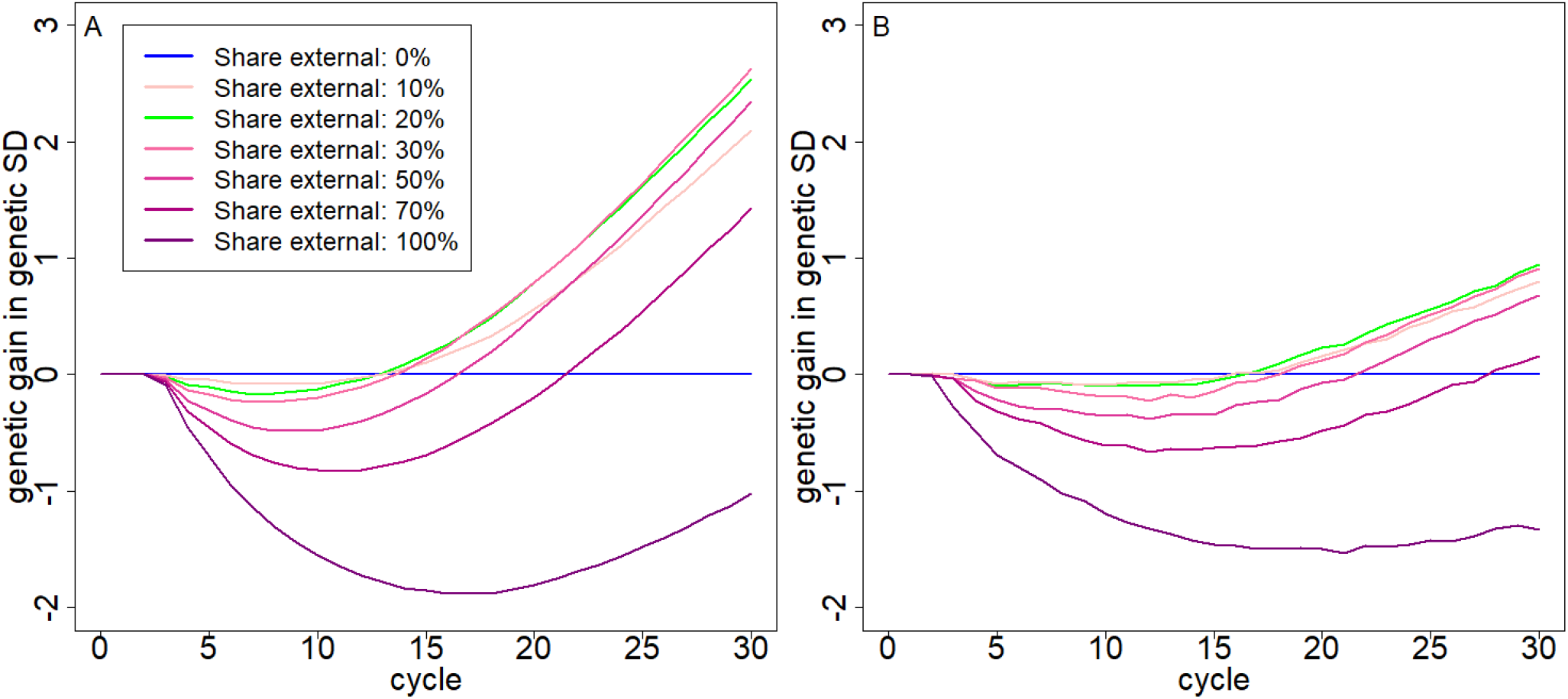
Genetic gain compared to the scenario with no newly introduced genetic material for the crosses (A) and DH lines (B) in the F2 lines of the respective cycle based on the used amount of genetic diversity from outside of the breeding program in each cycle.

### Additional phenotyping

When additional phenotyping of the F2 lines was performed, a strong increase in the performance as a cross (6.80 gSD; + 100%) and a small improvement as a DH line (4.54 gSD; + 6%) was observed. The obtained prediction accuracies for the different cycles are about 0.55 (Figure S6). Although gains were smaller for the DH line performance, the indirect selection based on the phenotypes in crosses was still more efficient than the use of the phenotypes of the founder DHs. When instead generating phenotypes for additional DH lines, the average genetic gains were 5.69 gSD (+67%) and 6.74 (+57%) for DH lines from top F2 lines with average prediction accuracies of 0.60 across the different cycles (Figure S6).

It is to note that additional phenotyping led to an even stronger drop in the remaining genetic diversity. The standard deviations of the genetic values among the F2 lines are reduced by about 87% after 30 cycles (Figure 3.C).

### Genetic diversity through introduction of new variation

When adding additional genetic material with similar genetic values to the current F2 population, this did not result in any genetic progress compared to the baseline scenario (Figure 3) with genetic gains of 3.37 / 4.10 gSD (- 1% / -4%). Genetic gains for both scenarios with additional phenotyping in the first few cycles were slightly worse because newly introduced genetic material was genetically slightly inferior. However, after around 15-20 cycles genetic gains were on par with the scenario without newly introduced genetic variance, with substantially higher gains in subsequent cycles. After 30 cycles genetic gains were 9.34 / 5.48 gSD (+174% / +28%) for the phenotyping of crosses and 6.65 / 7.78 gSD (+95% / +82%) for the phenotyping of additional DH lines for the cross / *per se* traits respectively. The genetic diversity in all three scenarios increased substantially to remain at around 60% of the genetic diversity of the initial breeding material with around 12% of all genotypes being heterozygous variants and only 35% of all markers being fixed after 30 cycles. The interested reader is referred to Supplementary Tables S1 & S2 for detailed numbers on the proportion of heterozygous variants and fixed markers for all scenarios and cycles.

Across replicates of the same simulation scenario, standard deviation ranging between 1.30 gSD (baseline) and 1.69 gSD (phenotyping of crosses and added diversity) were observed. As each scenario was simulated 250 times, the averages given here have a standard error of less than 0.1 for all scenarios, making all scenarios (except baseline vs. baseline with added diversity) statistically significantly different from each other (Two-sample t-test; (Student 1908)).

When varying the share of introduced material in each cycle, the highest gains were achieved when 20-30% of the generated F2s included material from outside the breeding pool. Short-term gains were minimally higher with a lower share of newly introduced genetic variation, while genetic gains dropped rapidly after 15 cycles. A higher share of added genetic diversity results in a lower short-term gain and similar per-cycle gains in later cycles. As one would expect, levels of genetic diversity increase with a higher share of newly introduced genetic variance (Figure S7).

### Genetic diversity through selection

All results on the impact of different selection intensities and mating allocation from AlphaMate are displayed relative to three basic scenarios: The baseline, additional phenotyping of crosses, and additional phenotyping of crosses with newly introduced genetic material.

#### AlphaMate

For the scenarios without introduction of new genetic material, the use of low angles in AlphaMate (Gorjanc *et al*. 2018) resulted in strict improvement compared to the respective basic scenario with both higher genetic gain (Figure 5) and greater remaining genetic diversity (Figure S8). Higher angles resulted in substantially higher genetic diversity, however, even after 30 cycles this did not translate into a higher genetic gain for an angle of 70. The optimal angle for the baseline scenario was 40 with a genetic gain of 3.70 gSD (+9%) and 50 for the phenotyping of crosses (8.71 gSD; +29%). When introducing additional genetic variance, relative genetic gains were low, the best performance being observed with an angle of 40 leading to 9.77 gSD (+4%).

**Figure 5.**
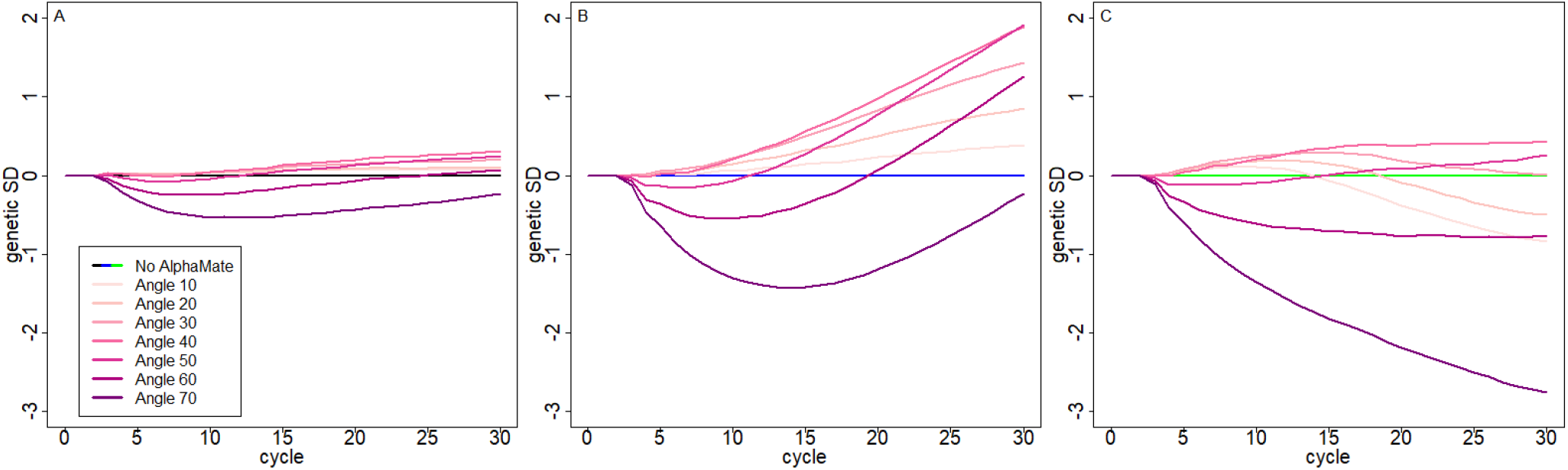
Genetic gain relative to the respective basic scenario based on the used angle between genetic gain and inbreeding in Alpha-Mate for the baseline scenario (A), a scenario with phenotyping of the DH lines (B), and phenotyped & introduction of genetic variance in each cycle of the rapid cycle breeding scheme (C). Coloring for reference scenarios is in concordance with Figure 3.

#### Selection intensity

When selecting fewer F2 lines per cycle, we observed reduced short-term gains compared to the respective basic scenario (Figure 6). However, after 5 to 7 cycles, the respective basic scenario already outperformed scenarios with 10 / 20 selected F2 lines with substantial drops in genetic gain (2.86 / 5.04 / 9.09 gSD; - 15 / -26 / -3%). Contrary, a lower selection intensity scenario outperforms the basic scenario after 10-15 cycles when no additional genetic variance was introduced with final genetic gains of 3.63 gSD (+7%) with no phenotyping and 8.45 gSD (+24%) with additional phenotyping of crosses after 30 cycles. The scenarios with newly introduced genetic material with selecting the top 20 / 30 / 50 F2 performed similarly to the basic scenario, while a substantially lower or higher number of F2 lines resulted in lower genetic gain (Figure 6).

**Figure 6.**
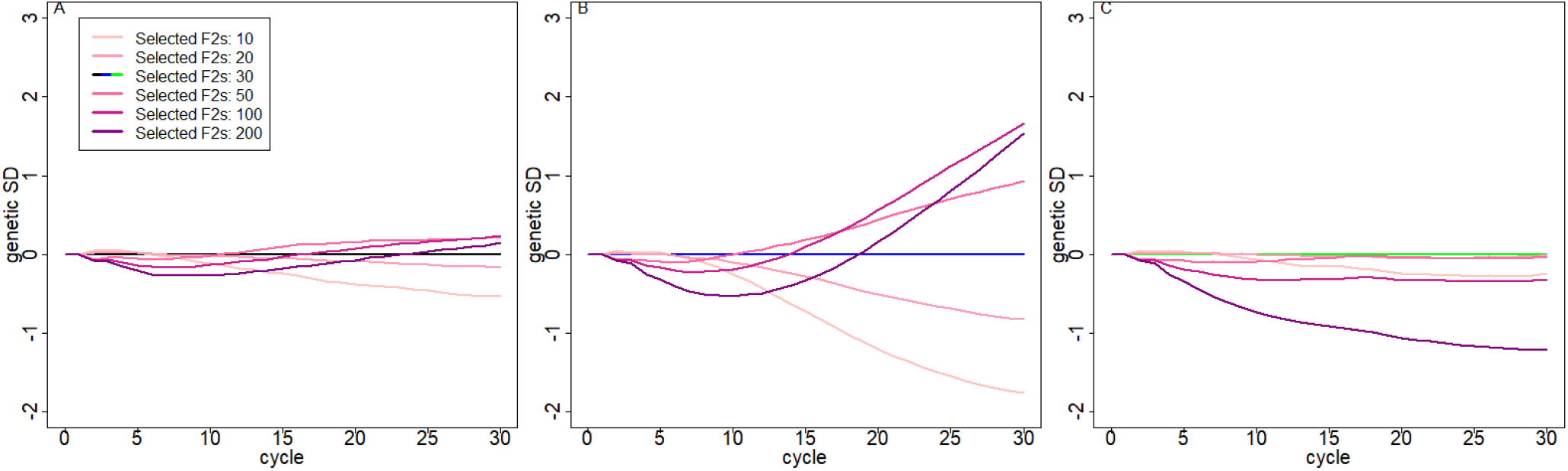
Genetic gain relative to the respective basic scenario depending on the selected F2 lines per cycle in the baseline scenario (A), additional phenotyping of crosses (B), and additional phenotyping of crosses with newly introduced genetic diversity (C).

When only 5 DH lines were used to generate the initial C1F1 lines, we observed substantially higher gains in the first breeding cycle (+0.1 gSD in all three basic scenarios; Figure 7). However, after just two breeding cycles, genetic gains were back on the same level as in the baseline scenario with 10 initially used DH lines, with substantially worse long-term performance for the two scenarios with no additionally introduced genetic material with gains of 3.15 / 6.29 gSD (−7 / -7 %). When adding no additional genetic diversity, the use of 50 founder DH lines resulted in longterm gains of 3.59 / 6.98 gSD respectively (+6 / +3%). When introducing genetic diversity from outside of the breeding pool, founding population sizes of 10, 20, 30, and 50 DH lines to generate the initial C1F1 led to very similar genetic gains.

**Figure 7.**
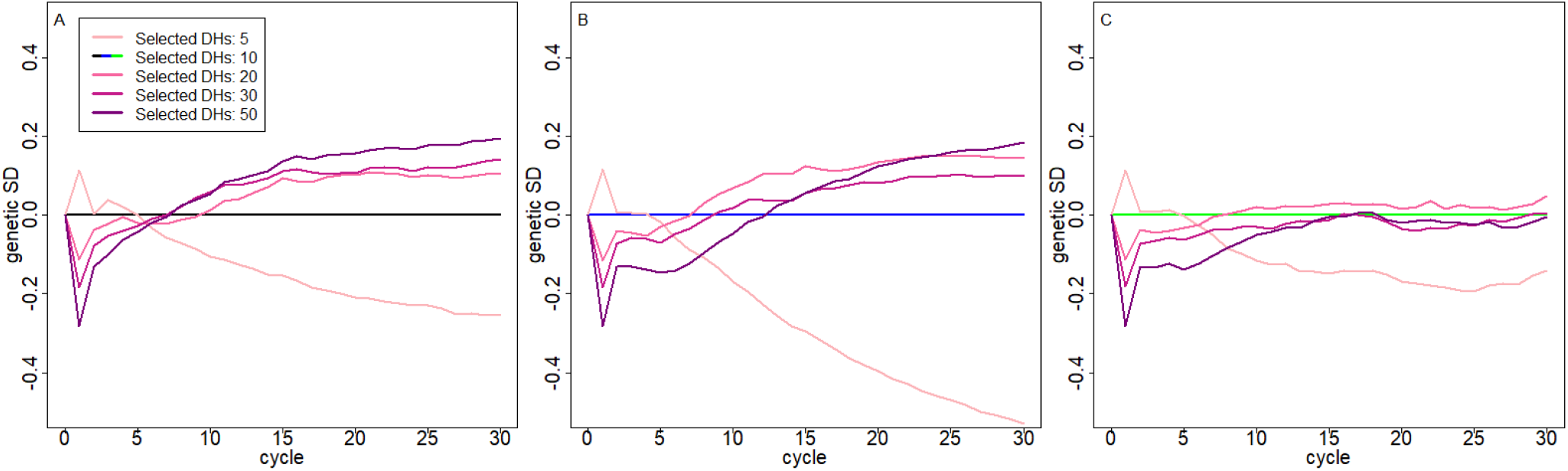
Genetic gain relative to the respective basic scenario depending on the number of initially selected DH lines to generate the C1F1 in the baseline scenario (A), additional phenotyping of crosses (B), and additional phenotyping of crosses with newly introduced genetic diversity (C).

### Computing / performance specs

The baseline scenario required 250 seconds of computing time of which 174 seconds were spent on the generation of new individuals and 72 seconds for breeding value estimation. Additional phenotyping increased computing times for the breeding value estimation to 192 seconds. Adding additional genetic diversity took an additional 12 minutes. All computations were performed using a single core on a system with a 2X Xeon Platinum 9242 3.80 GHz processor. Peak memory usage was between 2.5 to 3.3 GB for all scenarios.

Scenarios that used AlphaMate took 2.0 hours of runtime with a single run of AlphaMate requiring about 4.0 minutes.

## Discussion

This study shows via stochastic simulations that rapid cycle breeding as suggested by Polzer (2023) has the potential to be a valuable strategy in plant breeding to achieve substantial genetic gains in a limited time frame. However, to fully utilize its full long-term potential, proper management to ensure high prediction accuracies is of central importance. In the considered scenarios, prediction accuracies decrease after only a few breeding cycles and thus lead to significantly lower genetic gains. Thus, the generation of additional phenotypic data and the updating of the training model are of major importance. As correlations between *per se* and cross performance were high (0.6-0.8) phenotyping of both types of lines led to substantial improvements, with phenotyping of the crosses leading to higher gains for traits expressed as crosses and vice versa.

Depending on the breeding objective and time frame, the introduction of additional genetic diversity is of high importance to ensure long-term genetic progress (Hölker *et al*. 2019; Swarup *et al*. 2021). In case a rapid cycle scheme is “only” used for pre-breeding and to get new potential breeding material on an adequate level for introduction into an existing breeding scheme, this might not be necessary. For the here-considered maize rapid cycling scheme with genotypes from a European maize landrace, we observed higher or at least very similar genetic gains for the first 15 cycles of the scheme, without the introduction of new genetic material. This will of course be highly dependent on the species, initial genetic diversity, and selection intensity.

With regard to the used selection intensity, resulting from the number of initial used DH lines and the number of selected F2 lines per cycle, our study suggests that the efficiency of this strategy depends on whether additional genetic diversity was added to the breeding pool at later time points. When applying weaker selection, genetic gains were slightly lower for 5-10 cycles when no additional genetic material is added to the breeding pool (e.g. pre-breeding), while substantial long-term improvements were obtained. In breeding schemes for variety development, where new material is frequently added, the negative long-term effects of high selection pressure are much lower. Note here that factors like lethal mutations or passive selection due to lines not being able to produce DH lines or inbreeding depression were not considered in the simulations (Holland 2007; Sitonik *et al*. 2019; Zeitler *et al*. 2020).

In particular, when no additional genetic diversity was added to the breeding program, the use of AlphaMate (Gorjanc *et al*. 2018) was highly beneficial as it led to higher genetic gains and more remaining genetic diversity. Thus, it is a strict improvement of the breeding scheme without any disadvantages (besides minor computing costs). When using AlphaMate with an angle of about 40, we found the best overall performance. Slightly lower angles produced a larger short-term gain and slightly higher angles produced a larger long-term gain. A potential factor limiting the usefulness of AlphaMate in the scenario of additional phenotyping with introgressed material could be that AlphaMate does not account for the relatedness of the F2 lines within the breeding material to the introgressed material. Other approaches for mate allocation were not tested as they did not provide the required flexibility of equal contributions of all parents to the next cycle (Meuwissen 1997) or are not really designed for plant breeding applications with single-sex and their mating schemes (Kinghorn 2011; Wellmann 2017).

The design used in this study with three plant height traits is a simplification of reality, as plant height will usually not be the primary target trait. However, the results presented here should generalize to more sophisticated indices with more traits or non-linear effects. Here, all traits were considered polygenic, with predominantly purely additive and dominant QTLs and only a small amount of epistatic variation. No method for accounting for epistasis in the prediction model (Martini 2017; Vojgani *et al*. 2020) or specific breeding strategies to maintain beneficial epistatic QTLs were used (Holland 2001; Ali *et al*. 2020). Similarly, no fixed effect or genotype-by-environment interactions were simulated. We would assume that all scenarios should be impacted by such changes in a very similar way, thus not compromising the overall results of this study. Further studies are needed to analyze the impact of the here-considered breeding actions for traits with a higher share of epistatic effects (Khattab *et al*. 2010; Zaazaa *et al*. 2012).

This study represents on of the first published applications of MoBPS (Pook *et al*. 2020b) in plant breeding. Although the results of the simulation at first glance might not seem that surprising, we would argue that this should mostly be seen as a confirmation that the simulator is working in the intended way. Note that the added value of stochastic simulations does not solely lie in the result that a high selection intensity will lead to higher short-term genetic gains and lower long-term gains as the genetic diversity is strongly reduced early. Instead, stochastic simulations should be seen as a way to quantify the size of such effects without requiring assumptions on factors like the accuracy of the breeding value estimation, the remaining genetic diversity, or the effect of using a certain selection technique, as those will be implicitly calculated in the process of simulations (Hassanpour *et al*. 2023) and thus support breeders in the design of a breeding program. Although general observations of the results reported here should translate well to other breeding schemes and/or crops, the exact optima will be dependent on the population structure, trait architectures, and the design of the breeding scheme.

## Data Availability

Genotype data for the founder DH lines were taken from Mayer *et al*. (2020) with the data being processed as described in Pook *et al*. (2019). The data is publicly available at https://doi.org/10.6084/m9.figshare.12137142. Although the dataset contains 501,124 useable markers, this was down-sampled to reduce the number of markers to 50,000 and mimic a medium-density array.

The source code for all simulations and the MoBPS R-package can be found at https://github.com/tpook92/MoBPS. MoBPS version 1.9.06 was used for all simulations in this study.

The software AlphaMate is available at https://github.com/AlphaGenes/alphamate. AlphaMate version 0.2.0 was used for all simulations performed.

Supplementary files are available at FigShare:

Figure S1 provides a schematic overview of the rapid cycle breeding scheme with blue boxes indicating cohorts of lines with numbers indicating the number of lines included in a particular cohort. This visualization was generated via the MoBPS interactive environment from Pook *et al*. (2020a), Figure S2 with additional phenotyping of F2 lines, Figure S3 with additional phenotyping of DH lines and Figure S4 with newly introduced additional genetic material. Figure S5 provides information of the breeding value estimation for the F2 lines in the different breeding cycles for each of the three traits. Figure S6 provides an overview of per trait and cycle prediction accuracies for the different scenarios. Figure S7 provides a comparison of the remaining genetic diversity for the different base scenarios depending on the amount of genetic diversity from outside of the breeding program in each cycle. Figure S8 provides a comparison of the remaining genetic diversity for the different base scenarios depending on the angle used in AlphaMate (Gorjanc *et al*. 2018) in each cycle. Table S1 provides information on the share of heterozygous variants for all scenarios and cycles. Table S2 provides information on the share of fixed markers for all scenarios and breeding cycles.

## Acknowledgments

The authors thank project partners from the project MAZE for useful discussions and feedback.

## Literature cited

Ali M, Zhang L, DeLacy I, Arief V, Dieters M, Pfeiffer WH, Wang J, Li H. 2020. Modeling and simulation of recurrent phenotypic and genomic selections in plant breeding under the presence of epistasis. The Crop Journal. 8:866–877.

Allier A, Lehermeier C, Charcosset A, Moreau L, Teyssèdre S. 2019. Improving short and long term genetic gain by accounting for within family variance in optimal cross selection. Frontiers in genetics. 10:1006.

Allier A, Teyssèdre S, Lehermeier C, Moreau L, Charcosset A. 2020. Optimized breeding strategies to harness genetic resources with different performance levels. BMC Genomics. 21:1–16.

am Niehoff T, ten Napel J, Bijma P, Pook T, Wientjes YCJ, Hegedűs B, Calus MPL. 2024. Improving selection decisions with mating information by accounting for mendelian sampling variances looking two generations ahead. Genetics Selection Evolution. 56:41.

Bijma P, Wientjes YCJ, Calus MPL. 2020. Breeding top genotypes and accelerating response to recurrent selection by selecting parents with greater gametic variance. Genetics. 214:91–107.

Bonk S, Reichelt M, Teuscher F, Segelke D, Reinsch N. 2016. Mendelian sampling covariability of marker effects and genetic values. Genetics Selection Evolution. 48:1–11.

Büttgen L, Geibel J, Simianer H, Pook T. 2020. Simulation study on the integration of health traits in horse breeding programs. Animals. 10:1153.

Chaikam V, Mahuku G, Prasanna BM. 2012. Chromosome doubling of maternal haploids. Doubled haploid technology in maize breeding: theory and practice. pp. 24–29.

Crossa J, Pérez-Rodríguez P, Cuevas J, Montesinos-López O, Jarquín D, de Los Campos G, Burgueño J, González-Camacho JM, Pérez-Elizalde S, Beyene Y. 2017. Genomic selection in plant breeding: methods, models, and perspectives. Trends in plant science. 22:961–975.

Dreisigacker S, Pérez-Rodríguez P, Crespo-Herrera L, Bentley AR, Crossa J. 2023. Results from rapid-cycle recurrent genomic selection in spring bread wheat. G3: Genes, Genomes, Genetics. 13:jkad025.

Duvick DN. 2002. in professional plant breeding. Farmers, Scientists, and Plant Breeding: Integrating Knowledge and Practice. p. 189.

Endelman JB. 2011. Ridge regression and other kernels for genomic selection with r package rrblup. The Plant Genome. 4:250–255.

Falconer DS, Mackay TF. 1996. Introduction to quantitative genetics. Essex, England. .

Foley JA, Ramankutty N, Brauman KA, Cassidy ES, Gerber JS, Johnston M, Mueller ND, O’Connell C, Ray DK, West PC. 2011. Solutions for a cultivated planet. Nature. 478:337.

Gaikpa DS, Kessel B, Presterl T, Ouzunova M, Galiano-Carneiro AL, Mayer M, Melchinger AE, Schön CC, Miedaner T. 2021. Exploiting genetic diversity in two european maize landraces for improving gibberella ear rot resistance using genomic tools. Theoretical and Applied Genetics. 134:793–805.

Gaynor RC, Gorjanc G, Hickey JM. 2021. Alphasimr: an r package for breeding program simulations. G3: Genes, Genomes, Genetics. 11:jkaa017.

Gerald N, Frei UK, Lübberstedt T. 2013. Accelerating plant breeding. Trends in plant science. 18:667–672.

Ghosh S, Watson A, Gonzalez-Navarro OE, Ramirez-Gonzalez RH, Yanes L, Mendoza-Suárez M, Simmonds J, Wells R, Rayner T, Green P. 2018. Speed breeding in growth chambers and glasshouses for crop breeding and model plant research. Nature protocols. 13:2944–2963.

Gorjanc G, Gaynor RC, Hickey JM. 2018. Optimal cross selection for long-term genetic gain in two-part programs with rapid recurrent genomic selection. Theoretical and Applied Genetics. 131:1953–1966.

Gorjanc G, Hickey JM. 2018. Alphamate: a program for optimizing selection, maintenance of diversity and mate allocation in breeding programs. Bioinformatics. 34:3408–3411.

Gorjanc G, Jenko J, Hearne SJ, Hickey JM. 2016. Initiating maize prebreeding programs using genomic selection to harness polygenic variation from landrace populations. BMC Genomics. 17:1–15.

Hassanpour A, Geibel J, Simianer H, Pook T. 2023. Optimization of breeding program design through stochastic simulation with kernel regression. G3: Genes, Genomes, Genetics. p. jkad217.

Heslot N, Jannink JL, Sorrells ME. 2015. Perspectives for genomic selection applications and research in plants. Crop Science. 55:1– 12.

Hölker AC, Mayer M, Presterl T, Bolduan T, Bauer E, Ordas B, Brauner PC, Ouzunova M, Melchinger AE, Schön CC. 2019. European maize landraces made accessible for plant breeding and genome-based studies. Theoretical and Applied Genetics. pp. 1–13.

Holland JB. 2001. Epistasis and plant breeding. Plant breeding reviews. 21:27–92.

Holland JB. 2007. Genetic architecture of complex traits in plants. Current opinion in plant biology. 10:156–161.

Jannink JL. 2010. Dynamics of long-term genomic selection. Genetics Selection Evolution. 42:1–11.

Khattab SA, Esmail RM, Al-Ansary AM. 2010. Genetical analysis of some quantitative traits in bread wheat (triticum aestivum l.). New York Science Journal. 3:152–157.

Kinghorn BP. 2011. An algorithm for efficient constrained mate selection. Genetics Selection Evolution. 43:1–9.

Lorenz AJ. 2013. Resource allocation for maximizing prediction accuracy and genetic gain of genomic selection in plant breeding: A simulation experiment. G3: Genes, Genomes, Genetics. 3:481– 491.

Louwaars NP. 2018. Plant breeding and diversity: A troubled relationship? Euphytica. 214:114.

Martin R, Pook T, Bennewitz J, Schmid M. 2023. Optimization strategies to adapt sheep breeding programs to pasture-based production environments: A simulation study. Animals. 13.

Martini JWR. 2017. Incorporating Interactions and Gene Annotation Data in Genomic Prediction. Ph.D. thesis. Georg-August-Universität Göttingen.

Mayer M, Hölker AC, González-Segovia E, Bauer E, Presterl T, Ouzunova M, Melchinger AE, Schön CC. 2020. Discovery of beneficial haplotypes for complex traits in maize landraces. Nature communications. 11:1–10.

Mayer M, Presterl T, Ouzunova M, Bauer E, Schoen CC. 2018. Representing allelic diversity of maize landraces by libraries of doubled-haploid lines: paper # 97. German Plant Breeding Conference, Wernigerode, Germany. p. 138.

Melchinger AE, Brauner PC, Böhm J, Schipprack W. 2016. In vivo haploid induction in maize: comparison of different testing regimes for measuring haploid induction rates. Crop Science. 56:1127–1135.

Meuwissen THE. 1997. Maximizing the response of selection with a predefined rate of inbreeding. Journal of animal science. 75:934– 940.

Meuwissen THE, Hayes BJ, Goddard ME. 2001. Prediction of total genetic value using genome-wide dense marker maps. Genetics. 157:1819–1829.

Neyhart JL, Tiede T, Lorenz AJ, Smith KP. 2017. Evaluating methods of updating training data in long-term genomewide selection. G3: Genes, Genomes, Genetics. 7:1499–1510.

Polzer C. 2023. A rapid cycling selection experiment in maize landraces. Gordon Research Conference on Quantitative Genetics and Genomics. .

Pook T, Büttgen L, Ganesan A, Ha NT, Simianer H. 2020a. Mobpsweb: A web-based framework to simulate and compare breeding programs. bioRxiv. p. 2020.07.08.193227.

Pook T, Büttgen L, Ganesan A, Ha NT, Simianer H. 2021. Mobpsweb: A web-based framework to simulate and compare breeding programs. G3: Genes, Genomes, Genetics. 11.

Pook T, Freudenthal J, Korte A, Simianer H. 2020b. Using local convolutional neural networks for genomic prediction. Frontiers in genetics. 11:1366.

Pook T, Heise J, Herzog S, Simianer H. 2019. Deep learning - an alternative for genomic prediction? Quantitative Genetics and Genomics Gordon Research Conference. .

Sallam AH, Endelman JB, Jannink JL, Smith KP. 2015. Assessing genomic selection prediction accuracy in a dynamic barley breeding population. The Plant Genome. 8:plantgenome2014.05.0020.

Sargolzaei M, Schenkel FS. 2009. Qmsim: A large-scale genome simulator for livestock. Bioinformatics. 25:680–681.

Schaeffer LR. 2006. Strategy for applying genome–wide selection in dairy cattle. Journal of Animal Breeding and Genetics. 123:218– 223.

Schopp P, Müller D, Technow F, Melchinger AE. 2017. Accuracy of genomic prediction in synthetic populations depending on the number of parents, relatedness, and ancestral linkage disequilibrium. Genetics. 205:441–454.

Sitonik C, Suresh LM, Beyene Y, Olsen MS, Makumbi D, Oliver K, Das B, Bright JM, Mugo S, Crossa J. 2019. Genetic architecture of maize chlorotic mottle virus and maize lethal necrosis through gwas, linkage analysis and genomic prediction in tropical maize germplasm. Theoretical and Applied Genetics. 132:2381–2399.

Smith S, Bubeck D, Nelson B, Stanek J, Gerke J. 2015. Genetic diversity and modern plant breeding, In:, Springer. volume 7. pp. 55–88.

Student. 1908. The probable error of a mean. Biometrika. pp. 1–25.

Swarup S, Cargill EJ, Crosby K, Flagel L, Kniskern J, Glenn KC. 2021. Genetic diversity is indispensable for plant breeding to improve crops. Crop Science. 61:839–852.

Villiers K, Dinglasan E, Hayes BJ, Voss-Fels KP. 2022. genomicsimulation: fast r functions for stochastic simulation of breeding programs. G3: Genes, Genomes, Genetics. 12:jkac216.

Vojgani E, Pook T, Hölker AC, Mayer M, Schön CC, Simianer H. 2020. Bivariate genomic prediction of phenotypes by selecting epistatic interactions across years. bioRxiv. p. 2020.11.18.388330.

Voss-Fels KP, Stahl A, Wittkop B, Lichthardt C, Nagler S, Rose T, Chen TW, Zetzsche H, Seddig S, Baig MM. 2019. Breeding improves wheat productivity under contrasting agrochemical input levels. Nature plants. 5:706–714.

Wanga MA, Shimelis H, Mashilo J, Laing MD. 2021. Opportunities and challenges of speed breeding: A review. Plant Breeding. 140:185–194.

Watson A, Ghosh S, Williams MJ, Cuddy WS, Simmonds J, Rey MD, Hatta MAM, Hinchliffe A, Steed A, Reynolds D. 2018. Speed breeding is a powerful tool to accelerate crop research and breeding. Nature plants. 4:23–29.

Wellmann R. 2017. optisel: optimum contribution selection and population genetics. R Package. version 0.9.1. .

Zaazaa EI, Hager MA, El-Hashash EF. 2012. Genetical analysis of some quantitative traits in wheat using six parameters genetic model. American-Eurasian J. Agric. & Environ. Sci. 12:456–462.

Zeitler L, Ross-Ibarra J, Stetter MG. 2020. Selective loss of diversity in doubled-haploid lines from european maize landraces. G3: 12 Rapid cycle breeding in MoBPS Genes, Genomes, Genetics. 10:2497–2506.

